# Maternal expression of the novel centrosome assembly factor Wdr8 is required for vertebrate embryonic mitosis and development

**DOI:** 10.1101/031336

**Authors:** Daigo Inoue, Manuel Stemmer, Thomas Thumberger, Joachim Wittbrodt, Oliver J. Gruss

**Affiliations:** Centre for Organismal Studies (COS), Im Neuenheimer Feld 230, 69120 Heidelberg, Germany.; Zentrum für Molekulare Biologie der Universität Heidelberg (ZMBH), DKFZ-ZMBH Alliance, Im Neuenheimer Feld 282, 69120 Heidelberg, Germany.

## Abstract

The assembly of the first centrosome occurs upon fertilisation when the male centrioles recruit pericentriolar material (PCM) from the egg cytoplasm. While inaccuracy in the assembly of centrosomes leads to infertility and abnormal embryogenesis, the mechanism that ensures accurate assembly in vertebrate embryos remains obscure. Here we applied a CRISPR-Cas9-mediated gene knockout to show that Wdr8, a novel centrosomal protein, is maternally essential for PCM assembly during embryonic mitoses of medaka (*Oryzias latipes*). Maternal/zygotic Wdr8-null (Wdr8^−/−^) blastomeres exhibit severe defects in PCM assembly that cause them to divide asymmetrically and develop multipolar mitotic spindles and aneuploidy. We demonstrate that Wdr8 interacts via its WD40 domains with the centriolar satellite protein SSX2IP. Strikingly, exogenously provided Wdr8 fully rescues Wdr8^−/−^ embryos to adulthood, except in variants with mutations in the WD40 domains. This combination of targeted gene inactivation and *in vivo* reconstitution of the maternally essential Wdr8-SSX2IP complex reveals an essential link between maternal PCM and the stability of the zygotic genome in the early vertebrate embryo.

## Introduction

The animal microtubule organising centre (MTOC), or centrosome, comprises a pair of centrioles embedded in pericentriolar material (PCM)^1–4^. The centrosome duplicates during every cell cycle to define the two MTOCs of the bipolar mitotic spindle^2, 3^. An abnormal assembly or number of centrosomes can cause cell division failure, genomic instability and tumorigenesis in somatic cells^5–7^. Their accurate assembly, on the other hand, governs the progression of early cleavages in animal zygotes. Most animal oocytes eliminate centrosomes during oogenesis, supposedly to avoid centriole aging, parthenogenesis or abnormal embryonic mitoses with an excess number of centrosomes^3, 8, 9^. The developing oocytes accumulate maternal centrosomal factors to establish the initial zygotic centrosome upon fertilisation by assembling PCM around a paternally introduced centriole^8, 9^. This serves as a template for the numerous centrosomes that are created during subsequent cleavages. Ultimately, therefore, the success of fertilisation and embryonic development depends on the accurate assembly of the initial centrosome from paternal and maternal material during fertilisation^9^.

In somatic cells in culture, bipolar mitotic spindle and metaphase progression can occur normally in the absence of the centrosome^10, 11^. In contrast, it has been suggested that centrosomes in the *Drosophila, C.elegans* and sea urchin zygotes are essential for embryonic mitoses (i.e. cleavages)^12–16^. For example in *Drosophila,* a PCM protein Spd-2 and a centriolar protein Sas-4 appears to be largely dispensable for *somatic* mitosis in cultured *Drosophila* cells and later development (after midblastula transition) respectively, but is essential for embryonic mitoses via its centrosome assembly functions^15, 16, 17^ Therefore, embryonic centrosome assembly seems to require a special form of regulation involving specific maternal factors, but an analysis of the mechanisms and factors involved in this process have just begun. In contrast, surprisingly, such crucial role of the centrosome controlled by maternal centrosome factors in vertebrate embryos has been poorly understood. Undoubtedly, there is a compelling need to reveal the centrosomes’ physiological roles in vertebrate embyrogenesis to provide insights into development and infertility mechanisms in humans. However, it is still elusive partly due to the difficulty of peeling apart the distinct functions of maternal and zygotic factors using the knock out methods at hand.

However, a few zebrafish mutants have served as a basis for examining the physiological functions of maternal centrosomal factors in vertebrates^18–20^. The *cellular atoll* (*cea*) embryo, a maternal-effect Sas-6 mutant, exhibits a failure of centrosome duplication, and has thus revealed functions of centrosomal factors that have been conserved in human somatic cells and teleost embryos^19^. The lymphoid-restricted membrane protein (lrmp) mutant *futile* displays a failure of the attachment of centrosomes to pronuclei immediately after fertilisation, again demonstrating a maternal-specific function^20^. But many novel factors have scarcely been investigated in vertebrates, making it impossible to model the molecular networks required for centrosome assembly during the initial stages of embryogenesis and later development.

To gain molecular insights into the role of maternal-specific centrosomal factors in centrosome assembly *in vivo,* we used the vertebrate model medaka (*Oryzias latipes*)*^21^.* We analysed an uncharacterized WD40 repeat containing protein, Wdr8, (the orthologue of human WRAP73) that we recently identified in a screen for maternal proteins^22^. We applied CRISPR-Cas9-mediated targeted gene inactivation, and elicited specific centrosome assembly defects in the absence of maternal and zygotic Wdr8. This allowed us to unravel an essential role for Wdr8 in maternal PCM assembly. Subsequently we used mRNA injection, which mimics maternal gene expression, and observed a remarkable reconstitution of centrosome assembly, proper cell divisions and gross development until adulthood. This *in vivo* reconstitution strategy of maternal Wdr8 functions allowed us to perform a straightforward screening of mutant variants that revealed domains/modules of the Wdr8 protein essential for PCM assembly in living vertebrate embryos. Overall, this system clearly delivered molecular insights into Wdr8’s essential function in embryonic centrosome assembly.

## Maternal but not paternal Wdr8 is essential for symmetric cleavages of Medaka embryos

In a recent study we identified a number of proteins that were specifically upregulated in *Xenopus laevis* oocytes. Alongside the centrosome assembly factor SSX2IP^22^, our screen revealed an upregulation of Wdr8, a previously uncharacterized WD40 repeat-containing protein. Particularly interesting was the observation that the two proteins interacted with each other in *Xenopus* egg extracts (OJG, unpublished data).

To analyse Wdr8’s functions in early vertebrate development, we took advantage of the transparency of medaka embryos to carry out a cell biological analysis combined with a CRISPR-Cas9 genome editing approach to target and inactivate medaka Wdr8 (OlWdr8, hereafter referred to as Wdr8). This inactivation was achieved through the targeted integration of a GFP-stop cassette into exon-3 of the Wdr8 locus (Extended Data Fig. 1a, b). A cross of heterozygous parents yielded Wdr8’’ homozygous offspring that showed no obvious abnormalities during development and were phenotypically undistinguishable from wild-type fish through all stages to adulthood (Extended Data Fig. 2). However, when we compared the four combinations resulting from a cross of Wdr8^−/−^ and wild-type fish (Fig. 1a), we found a severe phenotype in all embryos from homozygous mothers. All maternal/zygotic Wdr8^−/−^ mutants (m/zWdr8^−/−^) exhibited cleavage cycles that were significantly delayed (by 20–30 min) following the first cleavage (Fig. 1b). The second division and the following cleavage cycles revealed abnormal asymmetric cleavages with a failure of cytokinesis (Fig. 1a), which were sometimes observed during the first division (Supplementary information (SI) Movie 3). In the absence of maternally provided Wdr8, these embryos failed to gastrulate or reach the neurula stage (St.18)^23^ (Fig. 1a). In contrast, the development of all embryos from homozygous fathers was indistinguishable from that wild-type embryos (Fig. 1a and Extended Data Fig. 3). These results demonstrate that maternally provided Wdr8 is essential for faithful cleavages in the large blastomeres of the early fish embryo.

**Figure 1.**
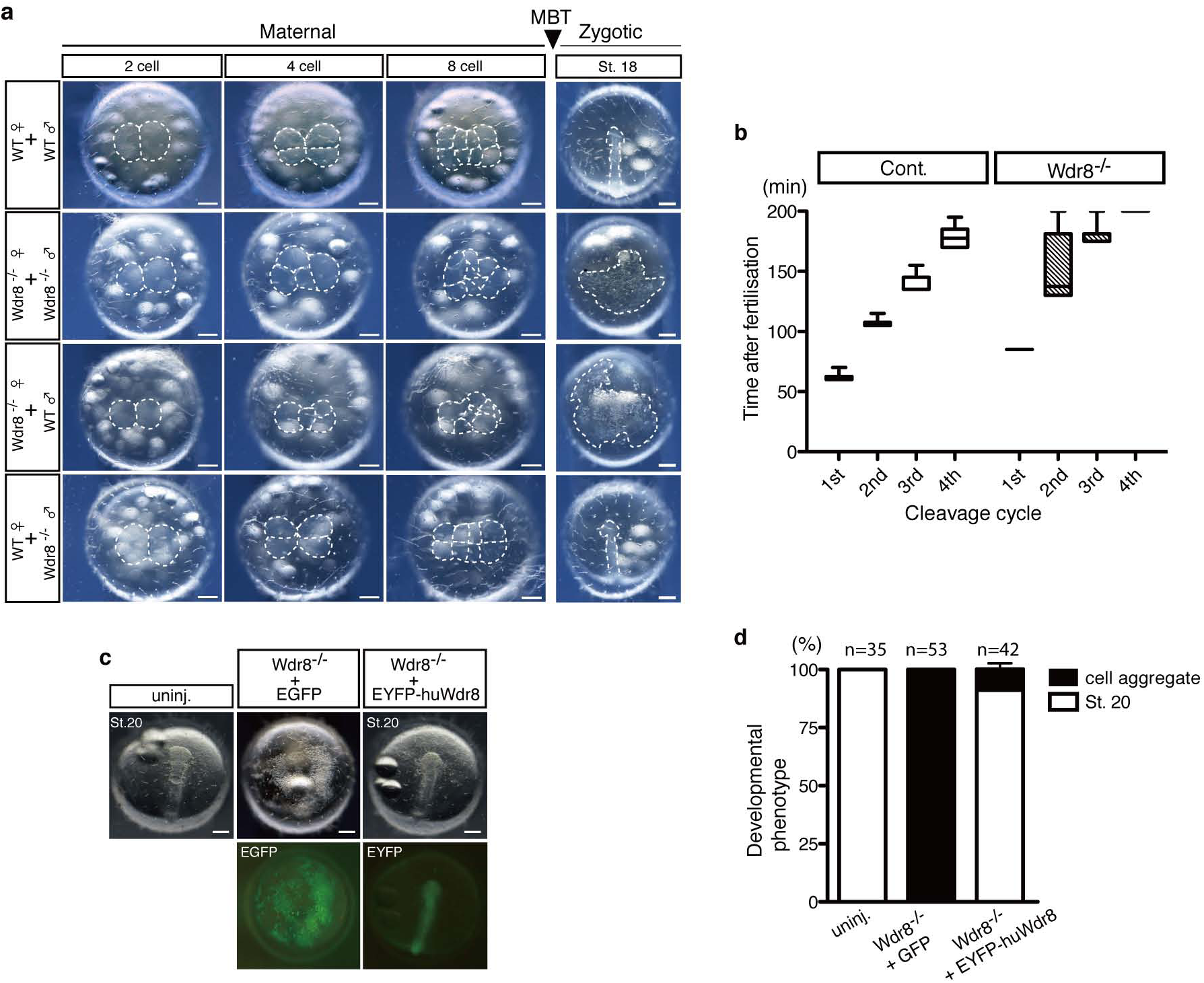
Abnormal cleavage divisions of Wdr8^−/−^ zygotes are rescued by EYFP-huWdr8 expression. **a,** Phenotypes of paternal and maternal Wdr8^−/−^ embryos before and after midblastula transition (MBT). The maternal Wdr8^−/−^ embryos, but not the paternal, showed disordered cleavage divisions and failure of gastrulation. **b**, Timing of cleavage cycles in Wdr8^−/−^ and wild-type specimens. Data represents median with minimum to maximum. Total embryos, 10 (Cont.), 11 (Wdr8^−/−^). **c, d**, Full rescue of Wdr8^−/−^ zygotes by exogenous expression of EYFP-huWdr8 (**c**) and its efficiency at St.20 (**d**). n, total embryos from three independent experiments. Data represent mean ± s.e.m. Scale bars, 200 μm.

## Exogenous expression of Wdr8 can efficiently rescue Wdr8^−/−^ zygotes

Given these indications that the presence of maternal Wdr8 during cleavage divisions suffices for the progression of normal development, we next performed rescue experiments in the maternal/zygotic mutants (Wdr8^−/−^). We injected mRNA encoding an EYFP-fusion of the human WDR8/WRAP73 orthologue (here called EYFP-huWdr8) into m/zWdr8^−/−^ zygotes within 5–10 min post fertilisation (mpf). At just 60 min after injection (about 70 mpf), functional proteins were detected from both the control mRNA, encoding EGFP alone, and EYFP-huWdr8. This was the point at which embryos underwent their first cleavage. While all the Wdr8^−/−^ embryos injected with EGFP showed abortive development, EYFP-huWdr8 expression fully rescued the cleavages and over 90 % of the injected eggs progressed through gastrulation to subsequent developmental stages (Fig. 1c, d). Intriguingly, about 70% of these embryos developed into juveniles indistinguishable from wild-type fish based on gross morphology (data not shown). These results not only confirmed the specificity of the knockout phenotype, but also clearly demonstrated that maternal Wdr8 plays an essential role in early development by maintaining the integrity of embryonic cleavages.

## Centrosomal localisation of Wdr8 in rescued Wdr8^−/−^ zygotes

The fluorescent tag of the rescue construct allowed us to determine the localisation of EYFP-huWdr8 in living blastomeres. *In vivo* microscopy showed that EYFP-huWdr8 localised specifically to one or two distinct dot-like domains in each blastomere of the rescued embryos (Fig. 2a, b). During the cleavage cycles from the one- to the four-cell stage, the two domains remained adjacent to each other, then separated and migrated to opposing sides of the blastomeres during mitosis (Fig. 2a and SI Movie 1). At the end of mitosis (before the emergence of the cleavage furrow), the Wdr8 domains were more dispersed (Fig. 2a and SI Movie 1). This localisation was strikingly reminiscent of the pattern that centrosomes follow during the cell cycle as they separate to form mitotic spindle. Strikingly, EYFP-huWdr8 co-localised with γ-tubulin as well as the centriolar satellite (CS)-marker PCM1^24,25^ at both prophase and metaphase, demonstrating that Wdr8 is a novel maternal centrosomal and CS factor in medaka embryos (Fig. 2c). In controls (EGFP-expressing wild-type and m/zWdr8^−/−^ blastomeres), the EGFP signal was found in the cytoplasm, confirming that the centrosomal localisation of the EYFP-huWdr8 signal is due to the Wdr8 sequence (SI Movies 1, 2, and 3). Taken together, our results indicate that Wdr8 acts as a centrosomal organiser in embryonic mitoses.

**Figure 2.**
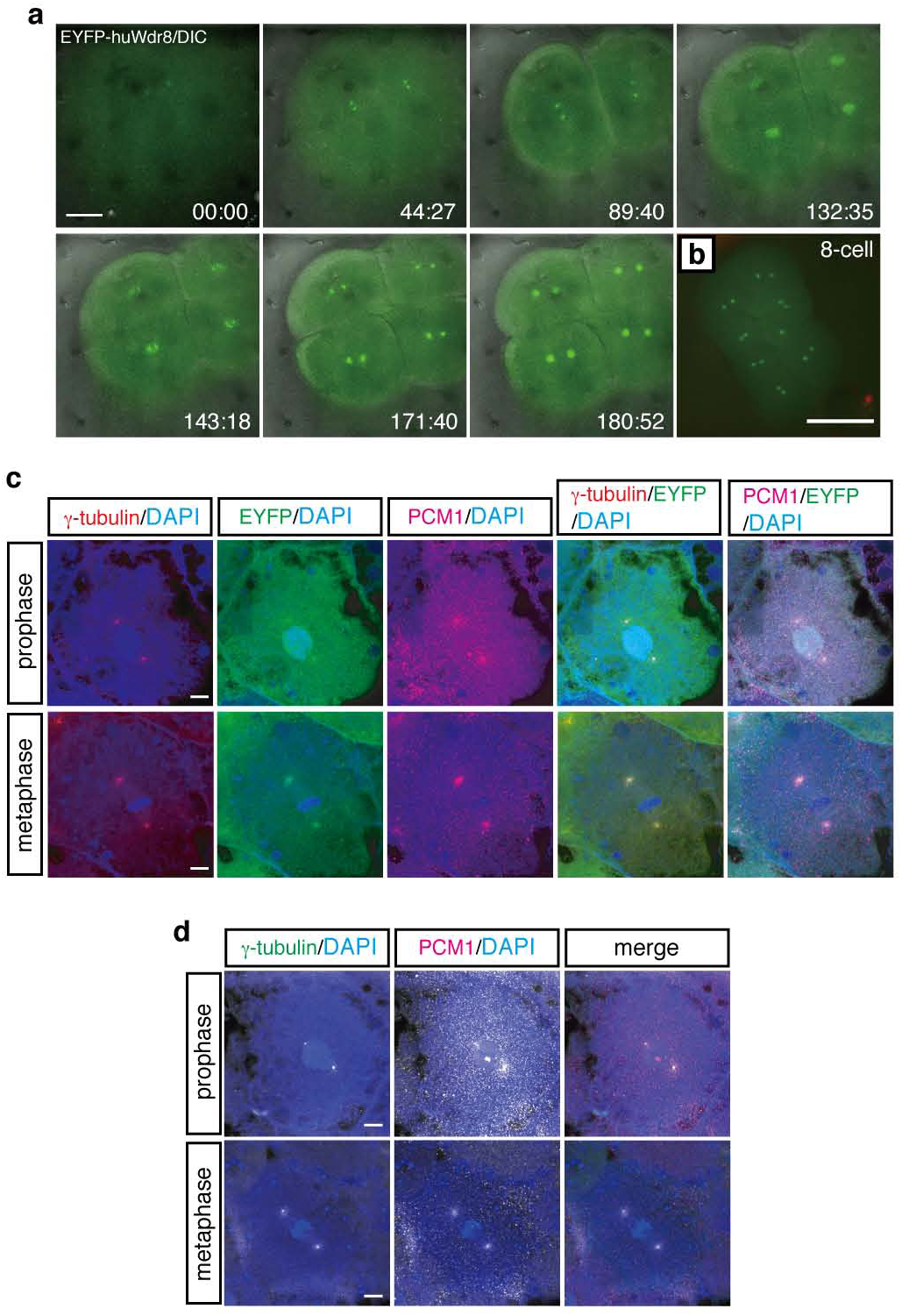
Wdr8 localises to the centrosome in the rescued Wdr8^−/−^ blastomeres. **a, b,** Spatiotemporal localisation of EYFP-huWdr8 in the rescued Wdr8’’ zygotes. Time-lapse images show stereotypic centrosome cycles from one- to 4-cell stage (**a**) and 8-cell stage (**b**). Time, min. **c, d,** Wdr8 is a novel centrosome protein. EYFP-huWdr8 colocalised with γ-tubulin and PCM1 (**c**), resembling the wild-type situation (**d**). Scale bars, 100 μm (**a**), 200 μm (**b**), 10 μm, (**c, d**).

## Wdr8 is essential for PCM assembly during rapid embryonic mitoses

The centrosomal localisation of γ-tubulin and PCM1 in the embryos rescued by EYFP-huWdr8 expression was comparable to that observed in wild-type embryos (Fig. 2d). This indicates that EYFP-huWdr8 successfully rescues the phenotype of Wdr8^−/−^ embryos by restoring the function of embryonic MTOCs. We therefore asked if MTOC assembly is affected in m/zWdr8^−/−^ blastomeres. At prophase and metaphase in these blastomeres, γ-tubulin and PCM1 were severely scattered compared to the wild-type, sometimes forming multiple PCM foci that were unevenly fragmented (Fig. 3a). Furthermore, while bipolar mitotic spindles readily formed in wild-type blastomeres, the scattered PCM of m/zWdr8^−/−^ blastomeres induced multipolar spindle assembly from prophase to telophase (Fig. 3b and Extended Data Fig. 4a, b). Importantly, these centrosomal and spindle abnormalities led to severe chromosomal instability (Fig. 3a, b and Extended Data Fig. 4a, b). In contrast, blastomeres expressing exogenous EYFP-huWdr8 recovered from all three defects: abnormal PCM assembly, multipolar spindle formation and aneuploidy (Fig. 2c and Fig. 3b). These results demonstrate that maternal Wdr8 ensures faithful PCM assembly and is essential for the maintenance of functional MTOCs in embryonic mitosis.

**Figure 3.**
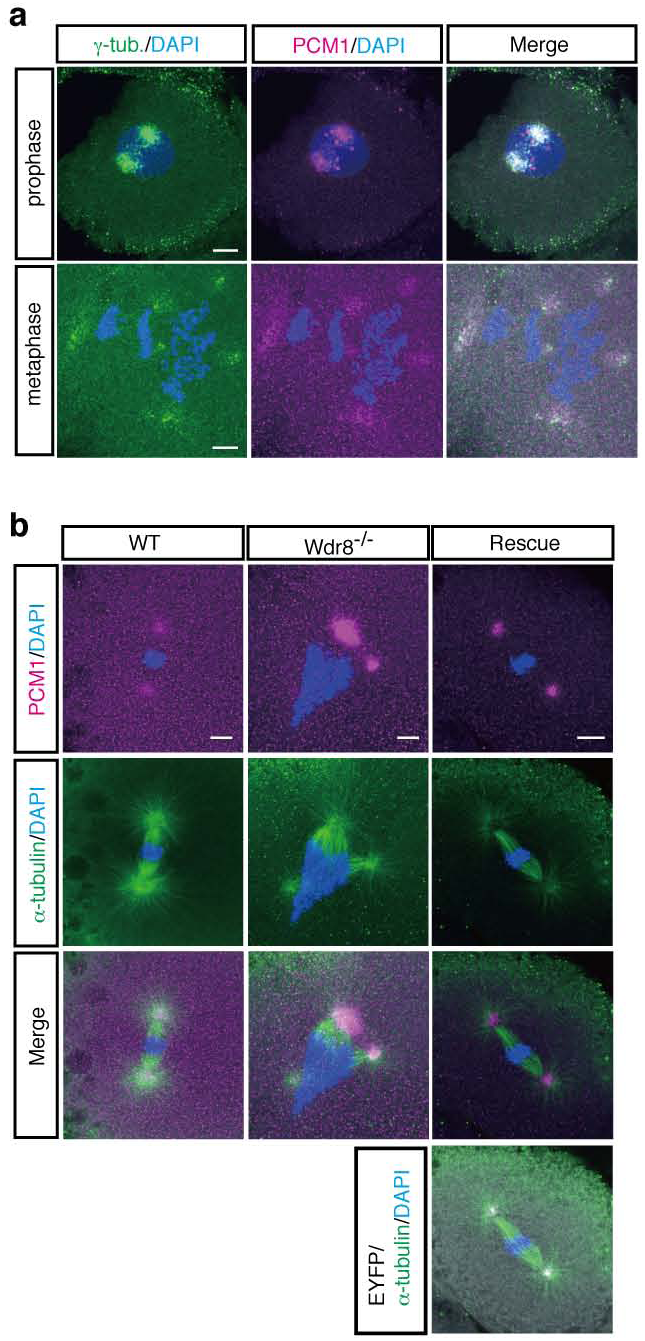
Centrosome and spindle abnormalities in Wdr8^−/−^ blastomeres. **a, b,** Abnormal PCM and mitotic spindle assemblies in Wdr8^−/−^ blastomeres. In comparison with wild-type (Fig. 2d), Wdr8^−/−^ eggs exhibited severely scattered γ-tubulin/PCM1 (**a**) and multipolar mitotic spindles (**b**) together with aneuploidy at metaphase (**a, b**). These abnormalities were fully rescued by EYFP-huWdr8 expression (Fig. 2c, Fig. 3b). Scale bars, 10 μm.

## The WD40 domains of Wdr8 are essential for both localisation and function of Wdr8

The fact that exogenously provided Wdr8 mRNA effects a highly efficient rescue during early cleavage stages provides a clear, systematic approach to the identification of interaction partners and the domains of Wdr8 that permit them to bind. Wdr8 is conserved across species from yeast to humans, and vertebrates exhibit four WD40 domains (Fig. 4a). WD40 domains are often involved in protein-protein interactions^26^, suggesting that they might play an important role in Wdr8’s function. We selected amino acids in the WD40 domains of Wdr8 that were conserved across species and mutated either W196D197 (WD40_1) or W359D360 (WD40_3) by Alanine (hereafter each mutant variant called 196/197AA and 359/360AA, Fig. 4a). To assess the functional relevance of the WD40 domain, we performed a rescue assay by injecting mRNA into m/zWdr8^−/−^ zygotes. mRNAs of all the variants (EYFP-huWdr8 wild-type, EYFP-196/197AA, or EYFP-359/360AA) were expressed at similar levels (Extended Data Fig. 5 and 359/360AA, data not shown). In contrast to the wild-type construct, Wdr8 lost its centrosomal localisation in both WD mutant variants and was mainly detected in the cytoplasm (Fig. 4b).

**Figure 4.**
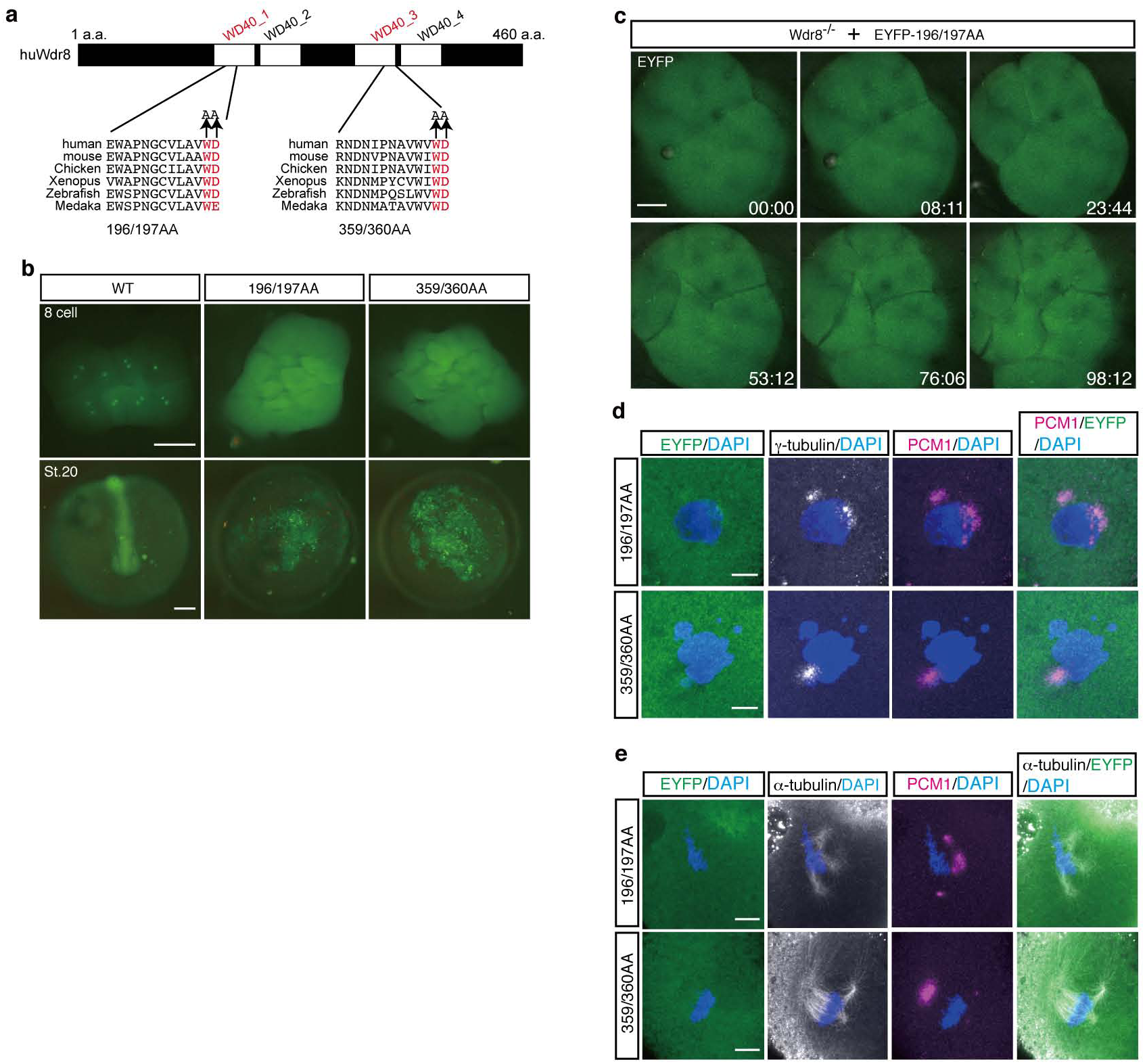
WD40 domains are essential for Wdr8 function. **a,** The conserved four WD40 domains in huWdr8. Mutations for WD mutant variants are denoted in red. **b,** External phenotypes of WD mutant-injected Wdr8^−/−^ embryos. Note that both variants mostly localised in the cytoplasm, unable to rescue Wdr8^−/−^ embryos. **c**, Time-lapse images of EYFP-196/197AA localisation in Wdr8^−/−^ blastomeres. EYFP-196/197AA weakly localised in the centrosome with multiple foci, which disappeared during cleavages. Time, min. **d, e**, Both mutant variants expression were unable to rescue abnormal PCM assembly (γ-tubulin/PCM1), multipolar mitotic spindles, and aneuploidy. Scale bars, 200 μm (**b**), 100 μm (**c**), 10 μm (**d, e**).

When blastomeres entered mitosis, a weak but visible centrosomal localisation of the mutant variants was observed in live imaging of either EYFP-196/197AA- or EYFP-359/360AA-injected embryos (Fig. 4c, SI Movie 4, time points at 08:11, 53:12, and 98:12). Individual blastomeres exhibited multiple foci, but they disappeared during the subsequent cleavage cycles (Fig. 4c, SI Movie 4). Interestingly, in m/zWdr8^−/−^ blastomeres, both WD variants were ubiquitously dispersed throughout the cytoplasm and slightly co-localised with scattered γ-tubulin and PCM1 at prophase (Fig. 4d). This indicates that WD variants are hypomorphic forms of Wdr8 that inefficiently localise to the centrosome, and demonstrates that the WD40 domain is essential for proper PCM/CS assembly.

Importantly, abnormal PCM/CS assembly led to multipolar spindles as well as chromosomal instability (Fig. 4e). This was consistent with the finding that neither of the WD mutant variants was able to rescue the early m/zWdr8^−/−^ phenotypes or their subsequent abortive development (Fig. 4b). In contrast, the mutation of four conserved putative Cdk1-phosphorylation motifs (mutations substituting AP for SP, a site S232 lies in WD40_2)^27^, affected neither the localisation nor the rescue function of Wdr8 in m/zWdr8^−/−^ embryos (data not shown). These results clearly demonstrate that intact WD40 domains govern the centrosomal localisation of Wdr8 and are required for proper PCM/CS functions in early embryonic mitoses of cleavage stages.

## The WD40 domains of the Wdr8-SSX2IP complex are essential specifically for its localisation and functions

We next took advantage of the mRNA-based *in vivo* reconstitution assay to address the molecular mechanisms underlying Wdr8’s function. We hypothesised that WD40 domains serve either as centrosome-targeting domains or as modules for interactions with other proteins. We first tested whether the rescue of m/zWdr8^−/−^ zygotes by WD variants was achieved by targeting them to the centrosome. We used the centrosome-targeting motif PACT to facilitate the accumulation of WD mutant variants at the centrosome^28^. Both PACT fusion constructs weakly localised to the centrosome in some embryos (Fig. 5a, PACT-359/360AA, data not shown). In these embryos, however, the embryonic lethality introduced by Wdr8 knockout was not rescued (1dpf in Fig. 5a, PACT-359/360AA, data not shown). This failure – even when Wdr8 was localised to the centrosome – strongly suggests that the WD40 domains of Wdr8 are not simply centrosome-targeting domains.

**Figure 5.**
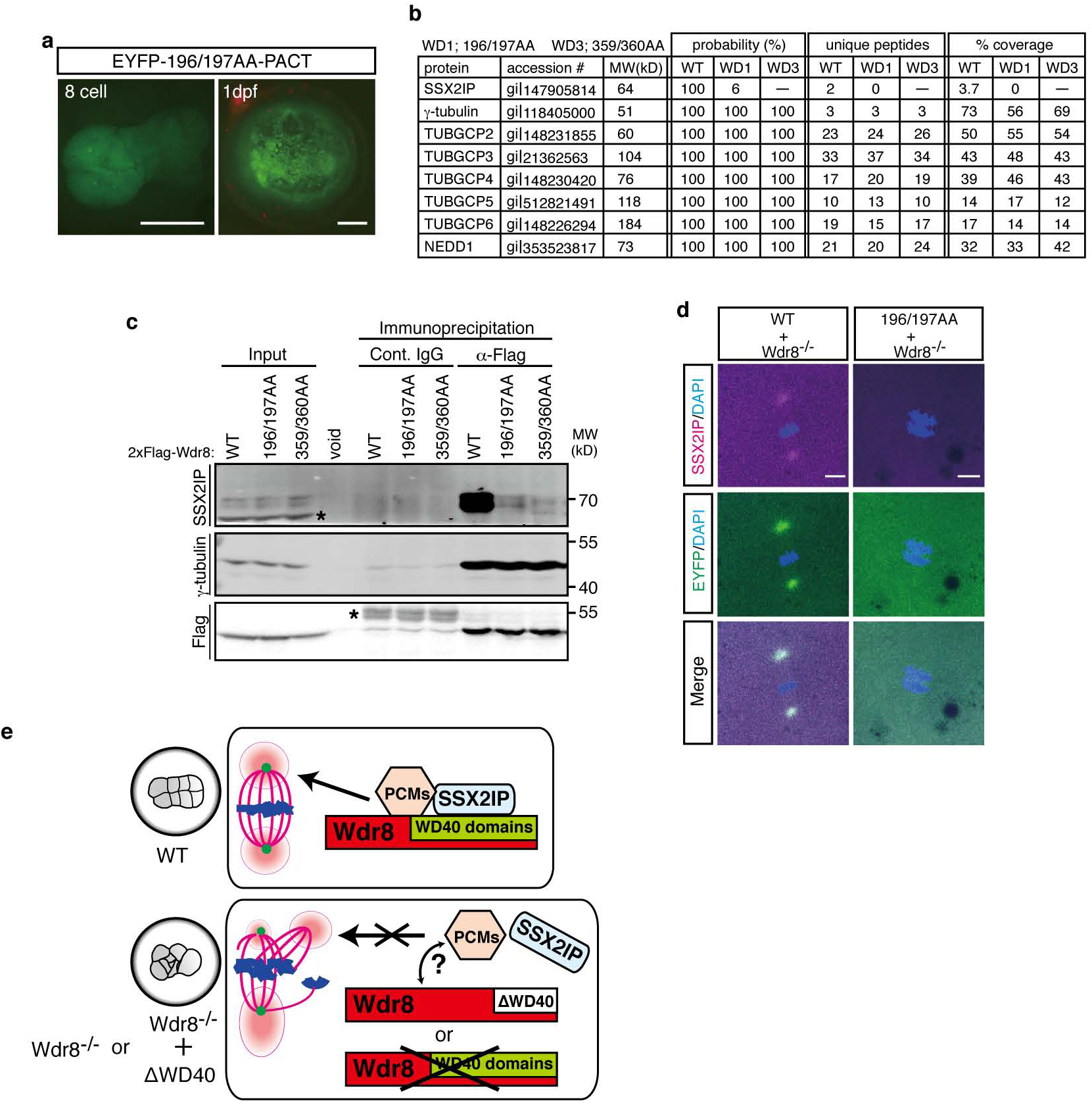
WD40 domains is essential to form Wdr8-SSX2IP complex and its localisation. **a,** Centrosome targeting of EYFP-196/197AA-PACT was insufficient for centrosomal localisation and the rescue of Wdr8^−/−^ embryos. **b, c,** Identification of Wdr8 interaction factors by mass spectrometry and immunoblot analyses with *Xenopus* egg extracts. WT, but not WD mutant variants, interacted with SSX2IP (**b, c**). Contrary, γ-TuRC subunits/Nedd1 (**b**) and γ-tubulin (**c**) interacted with Wdr8 independently of WD40 domains. *, background. **d**, Interdependent localisation of Wdr8 and SSX2IP via WD40 domains. Due to failure of Wdr8-SSX2IP interaction, EYFP-196/7AA, but not WT, was unable to colocalised with SSX2IP to the centrosome in Wdr8^−/−^ blastomeres. **e**, Wdr8-SSX2IP localises to the centrosome by tethering PCM components (e.g. γ-TuRC, which is essential for maternal PCM assembly). The absence of maternal Wdr8 or dysfunction of its WD40 domains causes PCM assembly defects. Scale bars, 200 μm (**a**), 20 μm (**d**).

Next we tested whether WD40 domains are important for interactions with other proteins. Since Wdr8 had initially been identified as an interaction partner of the CS protein SSX2IP, we hypothesised that WD40 domains might be critical for Wdr8’s interaction with SSX2IP or other centrosomal or CS proteins. To retrieve amounts of the proteins sufficient to analyse protein-protein interactions, we carried out a biochemical analysis of *Xenopus laevis* egg extracts. Following the expression of either 2xFlag-huWdr8 wild-type (WT), 2xFlag-196/197AA (196/197AA) or 2xFlag-359/360AA (359/360AA) in cytostatic factor (CSF) egg extracts arrested at metaphase^29^, we performed Flag immunoprecipitations to analyse the proteins that co-precipitated. Strikingly, mass spectrometry analysis revealed that SSX2IP interacted with WT, whereas it was unable to interact with either 196/197AA or 359/360AA (Fig. 5b), clearly demonstrating that WD40 domains are essential for the Wdr8-SSX2IP interactions.

Intriguingly, although γ-tubulin was severely scattered in Wdr8^−/−^ blastomeres, γ-tubulin and all the γ-TuRC subunits (including its regulatory protein Nedd1)^30–32^ interacted equivalently with WT and WD variants, indicating that WD40 domains are not required for Wdr8-γ-TuRC/Nedd1 interactions (Fig. 5b). Consistent with this finding, Western blot analysis confirmed that only WT, but not WD mutant variants, interacted with SSX2IP (Fig. 5c). On the other hand, γ-tubulin’s interaction with Wdr8 was independent of WD40 domains (Fig. 5c). Taken together, these results revealed that WD40 domains are the basis of the complex formed by Wdr8 with SSX2IP, but its interactions with γ-TuRC are independent of the domains. In medaka blastomeres, Wdr8 wild-type almost completely co-localised with SSX2IP, strongly suggesting that Wdr8 is a CS protein which forms a complex with SSX2IP (Fig. 5d). In contrast, SSX2IP was remarkably dispersed in the cytoplasm in both 196/197AA- and 359/360AA-expressing blastomeres (Fig. 5d, 359/360AA, data not shown), indicating that the interaction between Wdr8 and SSX2IP is crucial for mutual localisation and the function of Wdr8-SSX2IP as a CS protein complex (Fig. 5d, e).

In summary, these results clearly demonstrate that the Wdr8-SSX2IP complex is maternally essential for PCM assembly to ensure rapid embryonic mitoses.

## Discussion

Although it has long been suggested that centrosomes in vertebrate is absolutely essential for embryonic mitosis, the ultimate impact of centrosomes on vertebrate embryonic development and its molecular mechanism has been ill-defined. In this study, we for the first time reveal the detailed molecular mechanism of centrosome assembly specifically regulated by the novel maternal centrosomal protein Wdr8 during medaka embryonic mitoses (Fig. 5e). Centrosomes regulated by maternal Wdr8 are absolutely essential for mitotic spindle bipolarity and accurate inheritance of the zygote genome, which is distinct from a certain dispensability of centrosomes in somatic cells (Fig. 5e). Therefore, we here present a first major step toward solving the long-standing problems of physiological significance of centrosomes in early vertebrate development.

Several studies in vertebrate embryos have explored the regulation of the centrosome during embryonic mitoses^18–20, 33, 34^. A few zebrafish maternal-effect mutants, such as *cea* (Sas-6) or *futile* (Lrmp), have revealed functions in centrosomal regulation in the embryo^19, 20^. However, these studies failed to reveal further molecular insights due to the lack of precise gene targeting and the immense efforts of forward genetic screens for maternal genes in vertebrate models. Our CRISPR-Cas9 knockout medaka line clearly segregates the distinct functional contributions of maternal and zygotic Wdr8 during development. We show that early lethality of Wdr8^−/−^ embryos can be rescued with high efficiency either through maternal effects or the exogenous expression of a functional Wdr8 mRNA, which mimics the presence of a corresponding maternal mRNA. Strikingly, the rescued animals exhibited normal development and grew to become fertile adults. This reliable “reconstitution” rescue system, based on wild-type Wdr8 or several mutant variants, provides an unambiguous readout that directly links the molecular features of Wdr8 structure to its function and interactions with SSX2IP. This indicates that reverse genetic approaches with the CRISPR-Cas9 system are ideally suited to dissect maternal- and zygotic-specific aspects of centrosome regulation, which have been lacking for vertebrate systems.

Very recently, Wdr8 was shown to form a ternary complex with SSX2IP and the minus-end-directed kinesin Pkl1 (kinesin-14 homologue), which maintains minus-end pulling forces of microtubules (MTs) in fission yeast^35^. The SSX2IP-Wdr8-Pkl ternary complex itself is required for its localisation/function in the spindle pole body (SPB)^35^. The yeast complex provides a model case in which Wdr8-SSX2IP contributes to the generation of a pulling force by the MTs at the SPB, possibly by capping γ-tubulin^35, 36^. However, the question of whether Wdr8’s functions have been conserved through evolution has yet to be addressed. In cell culture, a Wdr8 partner, SSX2IP is shown to be required in CS for the recruitment of γ-tubulin ring complexes (γ-TuRCs) and other specific components into the PCM for the assembly of expanded mitotic PCM^22, 37^ While SSX2IP is not absolutely essential for the mitotic spindle bipolarity and metaphase progression in cultured cells^22, 37^, its function originally identified as maternal protein from *Xenopus* eggs has not been addressed. By unveiling the exclusive function of maternal Wdr8, our data show that Wdr8 interacts with SSX2IP via its WD40 domains and that this complex is essential for their mutual centrosome localisation/function in medaka embryonic mitosis. Although mass spectrometry analysis in our experiments could not identify kinesin-14 in either SSX2IP- or Wdr8-IP samples, the localisation of SSX2IP is dependent on dynein at least in *Xenopus* egg extracts^22^. Therefore, a complex of Wdr8-SSX2IP bound to dynein as a minus-end-directed motor could be a highly conserved molecular module whose function is crucial to ensure proper PCM assembly and maintain its structure during embryonic mitosis. Since SSX2IP is exclusive to CS^22, 37^, Wdr8-SSX2IP might be first recognised as a cargo by dynein at the CS granule to transport the complex, together with other PCM components, to the centrosome (Fig. 5e). The formation of the Wdr8-SSX2IP complex via WD40 domains may therefore be a critical step in the assembly of PCM components and the initiation of centrosome maturation (Fig. 5e)^38,39^. Intriguingly, our mass spectrometry analysis further demonstrates that Wdr8 also associates with all the γ-TuRC subunits, including its regulatory protein Nedd1, in a manner independent of WD40 domains (Fig. 5b). It is plausible that Wdr8 alone could serve as a platform for γ-TuRC/Nedd1 in either its transport to the centrosome or in stabilising the minus-end of MT.

Defects of the assembly of the centrosome after fertilisation have been directly linked to infertility and abnormal embryonic development in humans^40, 41^. So far the causes of infertility have mainly been attributed to the abnormalities of the sperm centriole^40, 41^. The effects of maternal centrosomal factors in oocytes have not yet received much attention. Considering that genomic instability in aging eggs causes infertility^42, 43^, it is possible that the downregulation or absence of maternal centrosomal factors during meiosis may contribute to infertility as well. Since Wdr8 is a critical maternal factor for genomic stability as well as centrosome assembly, the function of Wdr8 in relation to other interaction partners during meiosis needs to be addressed. On the other hand, the rescue of Wdr8^−/−^ embryos through either maternal effects or an introduction of exogenous Wdr8 expression causes them to undergo normal development, which suggests that zygotic Wdr8 is dispensable for later development (Extended Data Fig. 2, 3 and, Fig. 1a, c). It is possible, of course, that zygotic Wdr8 does have functions in later development beyond its role in PCM assembly, or their failure to produce a clear phenotype might be attributable to the presence of other, redundant factors.

## Methods

**Husbandry of fish and preparation of medaka zygotes.** Husbandry of fish and preparation of zygotes were routinely performed as described previously^44^. The husbandry and all the experiments were performed according to local animal welfare standards (Tierschutzgesetz §11, Abs. 1, Nr. 1, husbandry permit number 35–9185.64/BH Wittbrodt and mutagenesis permit number G-206/09) and in accordance with European Union animal welfare guidelines. The fish facility is under the supervision of the local representative of the animal welfare agency.

**Construction of CRISPR/Cas9 materials and generation of Wdr8 knockout medaka line.** With CCTop (http://crispr.cos.uni-heidelberg.de)^45^, a sgRNA was chosen to target exon-3 of *Oryzias latipes* Wdr8 (OlWdr8, NCBI reference sequence: XM_004070359.2) with least potential off-target sites in the remainder of the genome (Extended Data Table 1). Synthetic sgRNA oligos were annealed and cloned into pDR274 (addgene, #42250). Wdr8-sgRNA forward (5’-TAGGTCTCTCGAGCAGCCGGAC-3’) Wdr8-sgRNA reverse (5’-AAACGTCCGGCTGCTCGAGAGA-3’). The donor plasmid was created as described previously^45, 46^ via Golden GATEway cloning comprising the following sequences: 1) a specific sgRNA target site derived from GFP for *in vivo* linearisation of the donor vector (cf sgRNA-1 in ^45^), 2) a 665 bp homology flank corresponding to the upstream genomic sequence of the Wdr8-sgRNA target site, 3) a *GFP* variant (*GFP^var^*)*^46^* that was silently mutated to prevent CRISPR/Cas9-mediated cleavage by the donor-linearising sgRNA (see ^45^), 4) followed by a triple polyadenylation signal. The homology flank was cloned from Cab genomic DNA with primers: Forward (5’-GCCGGATCCATGGTCCTCAGACTCCCTGT-3’), Reverse (5’-GCCGGTACCCGGCTGCTCGAGTGACCACACCT-3’) to facilitate cloning into the Golden GATEway entry vector, the forward and reverse primers were extended with a BamHI or KpnI restriction site, respectively. To knock out OlWdr8, *GFP^var^* with stop codon was inserted by homology directed repair (HDR) to generate nonsense mutation, which resulted in the C-terminal deletion of roughly 500 amino acids, i.e. deletion of >75% of full-length OlWdr8. One-cell stage Medaka zygotes were co-injected with 10 ng/μl of donor plasmid, 15 ng/μl per sgRNA (Wdr8-sgRNA and sgRNA-1^45^), 150 ng/μl Cas9 mRNA diluted in nuclease free water. Founders were screened by genotype PCR (Extended Data Fig. 1b) with FinClip protocol^47^ since *GFP^var^* expression could hardly be detected, possibly due to impaired protein folding of *GFP* fused with N-terminal short fragment of OlWdr8. To achieve reproducible genotype PCR results, the program and the primer sets for the screening were fixed as below. The primers: for wild-type locus, forward (5’-AGTGTTCAAGCAGTCCAACCA-3’) and reverse (5’-TGAGGAGACTAGTCCAATTGAGC-3’), for *GFP^var^* insertion, forward (5’-AGTGTTCAAGCAGTCCAACCA-3’) and reverse (5’-GAACTTGTGGCCGTTTACGT-3’). The genotype PCR reaction with Taq polymerase (NEB): 1. 95°C for 1min, 2. 95 °C for 30s, 3. 65 °C for 30s, 4. 72 °C for 40s, repeated the cycle from step 2, 30 cycles. Heterozygous F1 fish were crossed with Cab. Heterozygous F2 fish were crossed with each other to generate F3 homozygous fish, which developed normally and grew to viable and fertile adults. By crossing F3 homozygous parents, the maternal effect of Wdr8^−/−^ homozygous zygotes (F4) was obtained to analyse the function of maternal Wdr8.

**Whole mount *in situ* hybridisation with medaka embryos.** For whole mount *in situ* hybridisation (WISH), a full-length OlWdr8 cDNA was isolated from cDNA library of wild-type embryos (St. 32)^23^, and then cloned into pGEM-T easy vector (Promega) to generate the antisense and the sense RNA probes. WISH was performed as described previously^44^. The pictures were taken under Nikon SMZ18 binocular microscope with the NIS-Elements F4.00.00 imaging software (Nikon).

**Preparation of cDNAs, mRNAs, and sgRNAs**. EYFP-huWdr8, 2xFlag-huWdr8, and their mutant variants were cloned into pCS2^+^ for mRNA generation. WD40 mutant variants, 196/197AA and 359/360AA, were generated by mutation PCR. To create PACT fusion construct, PACT domain (kindly provided by Elmar Schiebel, Zentrum für Molekulare Biologie der Universität Heidelberg (ZMBH), Heidelberg, Germany) was cloned and N-terminally fused immediate after a WD mutant variant by fusion PCR. mRNAs were transcribed with SP6 mMessegae mMachine kit (Life technologies). sgRNAs were transcribed with mMessage mMachine T7 Ultra kit (Life technologies).

**Whole mount fluorescent immunostainings with medaka blastomeres**. Whole mount fluorescent immunostainings were performed with 4hpf medaka blastomeres (St. 8) as described previously^44^, except for omission of the heating step, with anti-γ-tubulin (1:200 dilution, Sigma-Aldrich, T6557), anti-PCM1 (rabbit, 1:300 dilution, a gift from A. Merdes, Université de Toulouse, Toulouse, France), anti-*α*-tubulin (1:100 dilution, Sigma-Aldrich, T9026), and anti-GFP (1:300 dilution, Life technologies, A10262) antibodies. Secondary antibodies were Alexa Fluor 488 anti-mouse IgG (Life technologies), Alexa Fluor 488 anti-chicken IgY (Life technologies), Alexa Fluor 546 goat anti-mouse IgG (Life technologies), Dylight 549 goat anti-rabbit IgG (Jackson ImmunoResearch), and Alexa Fluor 647 goat anti-rabbit IgG (Life technologies). All secondary antibodies were incubated at 1:200 dilutions. DNA was counterstained with DAPI (1:200 dilution from 5mg/ml stock, Sigma-Aldrich, D9564). Images were taken with the confocal microscopy (Leica TCS SPE) with either 20x water (Leica ACS APO 20x/0.60 IMM CORR) or 40x oil (Leica ACS APO 40x/1.15 Oil CS 0.17/E, 0.27) objectives and processed with Image J. For SSX2IP detection, 100 ng/μl anti-xlSSX2IP antibody was injected into one-cell stage of medaka zygotes, and then subjected to immunostaining as described previously^22^.

**Live imaging of medaka embryos**. Fluorescence in injected medaka embryos was checked at 70–90 min post injection. The embryo with chorion was incubated in 3% methyl cellulose/1xERM with the glass bottom dishes (MaTeck corporation). Live imaging was performed by Leica SPE confocal microscope with 20x water objective. Images were taken every 37s-49s (Movie 1–4) with a 2–3 μm- z-step size. Maximum z-stack projection images were processed with Image J and then converted to Quick Time movie files.

**Rescue experiments by injection of either EYFP-huWdr8 mRNA or its individual mutant variants’ mRNA.** Wdr8^−/−^ fertilised eggs were immediately collected after checking mating, and injected with mRNA within 5–10 min post fertilisation in the precooled 0.5x ERM medium. 100 ng/μl of mRNA of either EYFP-huWdr8 or its individual mutant variants was injected into Wdr8^−/−^ zygotes at about 1/3 volume of one-cell stage. These zygotes were incubated in 0.5x ERM at 28 °C. Embryos at 4 hpf (St. 8) were fixed with 4% PFA in 1x PTw to perform whole mount fluorescent immunostainings. Images of external phenotypes of live embryos were taken under Nikon SMZ18 binocular microscope with the NIS-Elements F4.00.00 imaging software (Nikon).

**Western blot analysis of medaka embryos at morula stage**. 400 ng/μl of mRNA encoding either EYFP-huWdr8 or WD mutant variant was injected into one-cell stage zygotes. The embryos were incubated for 28 °C for 2h, and then treated with hatching enzyme for another 2h to remove chorion. 50 embryos (around St. 8) for each were softly homogenised with 100 μl cold PBS to remove the yolk. After centrifugation at 3,000 rpm for 2 min at 4 °C, the supernatant was removed and the pellet of embryos was frozen at liq N_2_ and stored at −80 °C. The pellet was lysed on ice with 25 μl RIPA buffer (50 mM Tris-HCl at pH 8.0, 150 mM NaCl, 5 mM EDTA, 15 mM MgCl_2_, 1% Triton-X100) containing 10 μM pepstatin, 10 μg/ml aprotinin, 0.1 mM PMSF, 1 mM Na_3_VO_4_, and 1 mM NaF. After centrifugation at 14,000 rpm for 5 min at 4 °C, 20 μl supernatant was mixed with the equal volume of 2x Laemmli sample buffer containing 10% 2-mercaptoethanol, and boiled for 10 min at 100 °C, subjected to SDS-PAGE.

**Immunoprecipitation of 2xFlag-huWdr8 from *Xenopus* CSF egg extracts**.

*Xenopus* CSF egg extracts were routinely prepared as previously described^48^. 80 μl of the extracts was incubated with 3 μg mRNA of either 2xFlag-huWdr8 wild-type, 2xFlag-196/197AA, or 2xFlag-359/360AA for 90 min at 23 °C to express Flag-tagged fusion proteins in the extracts. The immunoprecipitations were performed with 2 μg of either anti-Flag antibody (Sigma-Aldrich, F1804) or normal mouse serum (Invitrogen, #1410) for 45 min at 23 °C, followed by incubation with 20 μl of Protein G Sepharose 4 Fast Flow (GE Healthcare Life Sciences) slurry (in CSF-XB) for another 25 min at 23 °C by tapping every 5 min. The Protein G Sepharoses were washed twice and one time with 250 μl of TBS-T (10 mM Tris-HCl at pH7.5, 0.5% tween-20, 150 mM NaCl) and 250 μl of TBS (10 mM Tris-HCl at pH7.5, 150 mM NaCl), respectively. After removing TBS, the samples were prepared with 30 μl of 2x Laemmli sample buffer containing 10% 2-mercaptoethanol, boiled for 10 min at 100 °C, subjected to Western blot and mass spectrometry analyses as described previously^22, 49^.

**Dimethyl labeling and mass spectrometry**. For quantitative mass spectrometry analysis, comparison of control IgG pull-down and 2xFlag-huWdr8 pull-down samples were performed by after dimethyl-labelling using stable isotopes^49, 50^. Original mass spectrometry data were analysed using Proteome Discoverer 1.4 and Mascot (Matrix Science; version 2.4) as described previously^22, 49^. Identified proteins were displayed by Scaffold_4.4.3 (Proteome Software Inc.) to retrieve significantly interacted proteins with either Wdr8 wild-type or WD40 mutant variants.

**Supplementary information** is linked to the online version of the paper at www.nature.com/nature.

## Acknowledgements

We thank Elmar Schiebel and Peng Liu for providing constructs and *Xenopus* egg extracts. We also appreciate Gislene Pierre and Wenbo Wang for providing constructs and sharing unpublished results. D.I. was supported by the Human Frontier Science Program (HFSP) long-term fellowship and the Japan Science Promotion Society (JSPS) fellowship for research abroad. O.J.G. was recipient of a start-professorship of the German excellence initiative, ZUK 49/TP1–16, as part of ZUK 49: Institutional strategy to promote top level research awarded to Heidelberg University. This project was supported by the ERC Advanced Grant - Manipulating and Imaging Stem Cells at Work (J.W.).

## Author contributions

O.J.G. supervised the project. D.I. and O.J.G. conceived and designed the project with critical input from J.W. D.I. performed all the experiments and analysed the data. M.S. and T.T established CRISPR-Cas9 system. D.I. and O.J.G. wrote the manuscript. J.W. supervised and supported D.I.

## Author information

The authors declare no competing financial interests. Reprints and permissions information is available at www.nature.com/reprints. Correspondence and request for materials should be addressed to O.J.G. and D.I.

## Extended Data Figure and Supplementary information legends

**Extended Data Figure 1.**
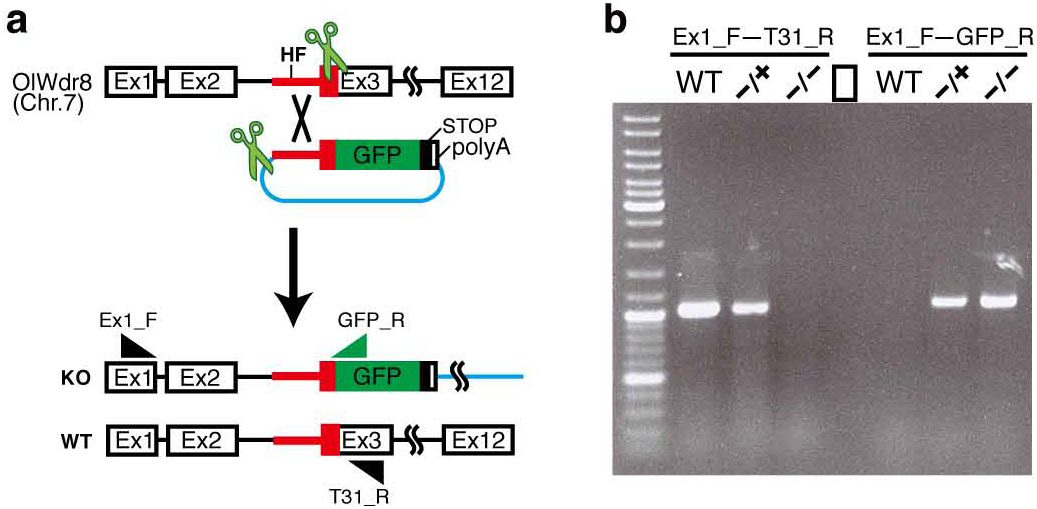
CRISPR-Cas9 mediated knockout strategy to generate Wdr8^−/−^ medaka line. **a,** A sgRNA was designed to knockout (KO) exon-3 of OlWdr8 by inserting 5’ homology flank (HF) and GFP with a stop codon (see details in Methods). **b,** The screening of Wdr8^−/−^ fish by genotype PCR. Two sets of primers, Ex1_F-T31_R and Ex1_F-GFP_R, were used to detect wild-type and the KO genomic locus, respectively. Note that WT locus was detected from genomic DNAs of WT and Wdr8^−/+^, but not that of Wdr8^−/−^. On the other hand, the KO locus was detected from genomic DNAs of Wdr8^−/+^ and Wdr8^−/−^, but not that of WT, due to the presence of GFP insertion.

**Extended Data Figure 2.**
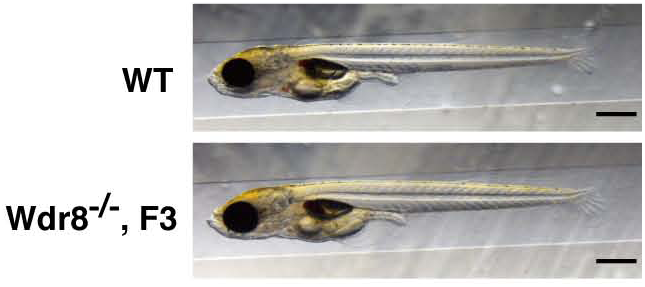
External phenotype of Cab wild-type and Wdr8^−/−^ hatchlings (F3 generation, see Methods). Wdr8^−/−^ hatchlings rescued by maternal Wdr8 developed normally with no obvious external abnormalities, comparable to wild-type. Scale bars, 500 μm.

**Extended Data Figure 3.**
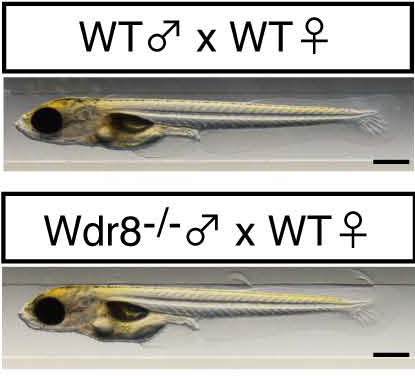
External phenotype of the hatchling from Wdr8^−/−^ father in comparison with wild-type hatchling. As in Extended Data Figure 2, there were no obvious differences, demonstrating that paternal Wdr8 is not implicated in both early and later stage development of medaka. Scale bars, 500 μm.

**Extended Data Figure 4.**
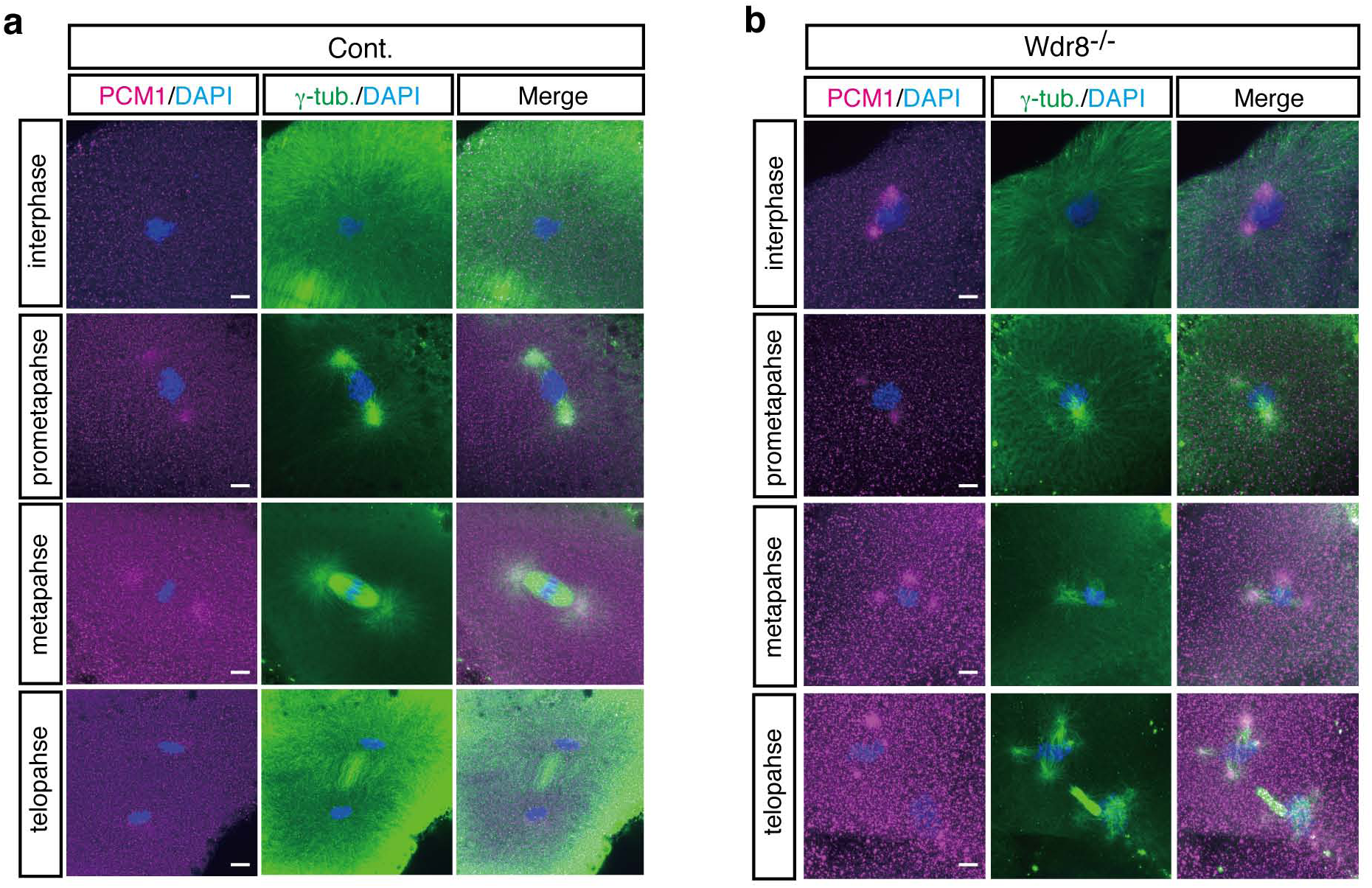
Detailed process of mitotic spindle assembly in Cab and Wdr8^−/−^ blastomeres. Whereas bipolar mitotic spindle was faithfully formed from prometaphase to telophase in wild-type blastomeres, fragmented centrosomes (PCM1) caused multipolar spindles as well as severe chromosome instability during mitosis in Wdr8^−/−^ blastomeres. Scale bars, 10 μm.

**Extended Data Figure 5.**
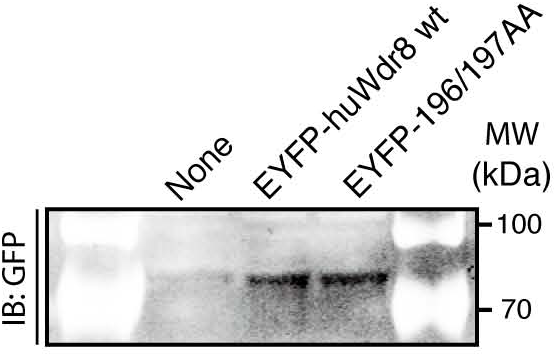
Western blot analysis to compare the protein level between EYFP-huWdr8 and EYFP-196/197AA. Expression levels of both proteins were almost equivalent, demonstrating that mutation in WD40 domain did not affect on protein stability.

**Extended Table 1.**
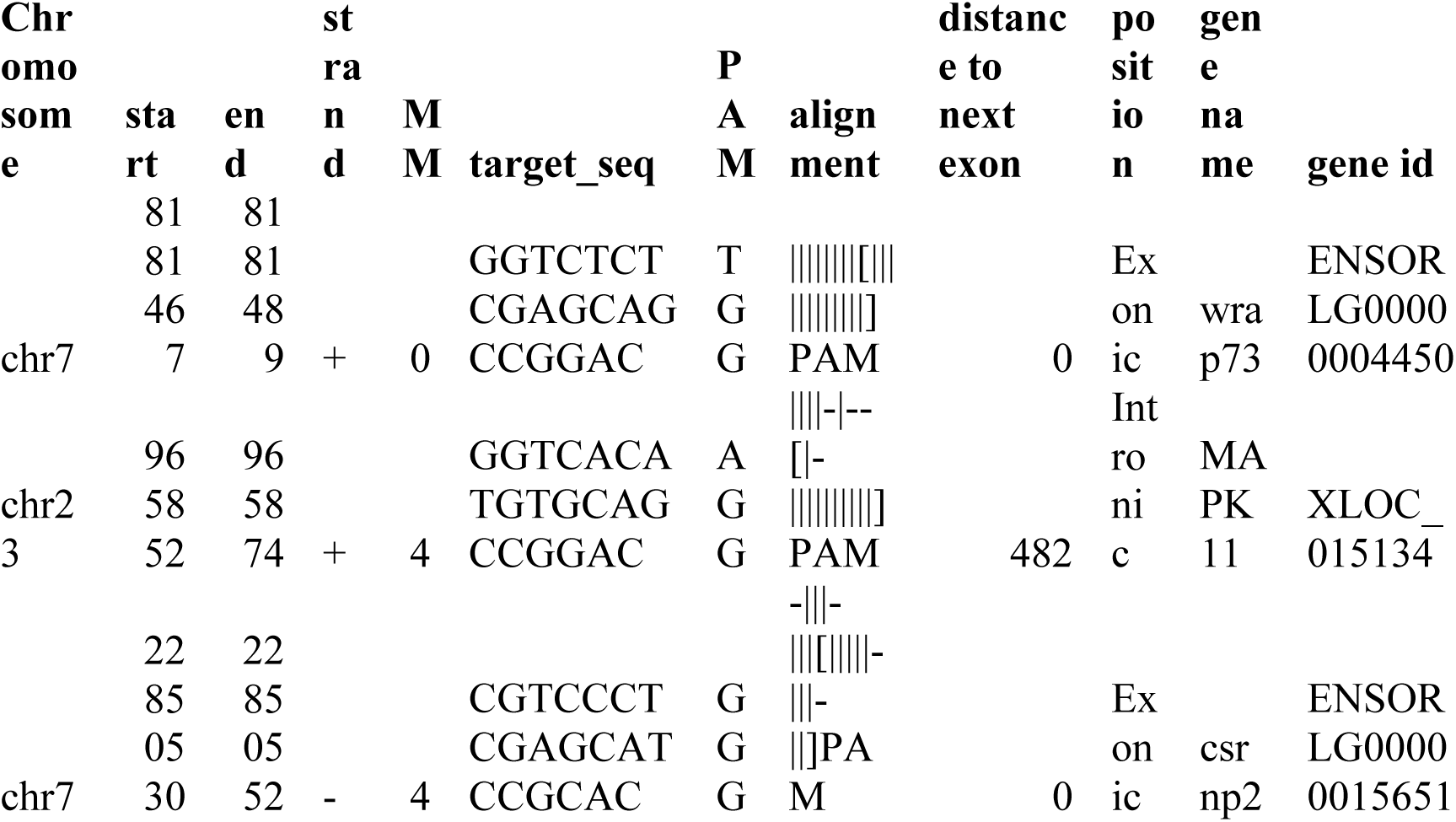
The chosen Wdr8-sgRNA target site has only two potential off-target sites in the entire medaka genome. Search-parameters (see ^40^ for further details): core length = 12, max. core mismatches = 2, max. total mismatches = 4, PAM = NGG

**SI Movie 1 Live imaging of EYFP-huWdr8 localisation in the rescued Wdr8^−/−^ embryos.** EYFP-huWdr8 localised to the one or two dots in individual blastomeres, showing stereotypic centrosome cycles during embryonic cell cycles from one- to 4- cell stage.

**SI Movie 2 Live imaging of EGFP localisation from one- to 4-cell stage of Cab wild-type embryos.** EGFP alone did not localise to the centrosome-like dots as EYFP-huWdr8 did in Wdr8^−/−^ blastomeres.

**SI Movie 3 Live imaging of EGFP localisation from one- to 4-cell stage of Wdr8^−/−^ embryos.** EGFP alone did not localise to the centrosome-like dots as EYFP-huWdr8 did in Wdr8^−/−^ blastomeres. Note that Wdr8^−/−^ disorderly divided blastomeres with cytokinesis failure.

**SI Movie 4 Live imaging of EYFP-196/197AA localisation from two-cell stage of Wdr8^−/−^ embryos.** EYFP-196/197AA mostly localised to the cytoplasm but oscillated weak centrosomal localisation with multiple foci during cleavages. Note that EYFP-196/197AA was not able to rescue the abnormal cleavage divisions of Wdr8^−/−^ embryos.

